# The Design of Brainstem Interfaces: Characterisation of Physiological Artefacts and Implications for Closed-loop Algorithms

**DOI:** 10.1101/2023.02.02.526892

**Authors:** Alceste Deli, Robert Toth, Mayela Zamora, Amir P. Divanbeighi Zand, Alexander L. Green, Timothy Denison

**Author notes:** This work was funded by the NIHR Oxford Biomedical Research Center and the Oxford Medical and Life Sciences Translational Fund. Tim Denison was funded by the Royal Academy of Engineering for device development. Correspondence: Timothy Denison for device-related queries, Alceste Deli for targeting and analyses-related queries.

## Abstract

Surgical neuromodulation through implantable devices allows for stimulation delivery to subcortical regions, crucial for symptom control in many debilitating neurological conditions. Novel closed-loop algorithms deliver therapy tailor-made to endogenous physiological activity, however rely on precise sensing of signals such as subcortical oscillations. The frequency of such intrinsic activity can vary depending on subcortical target nucleus, while factors such as regional anatomy may also contribute to variability in sensing signals. While artefact parameters have been explored in more ‘standard’ and commonly used targets (such as the basal ganglia, which are implanted in movement disorders), characterisation in novel candidate nuclei is still under investigation. One such important area is the brainstem, which contains nuclei crucial for arousal and autonomic regulation. The brainstem provides additional implantation targets for treatment indications in disorders of consciousness and sleep, yet poses distinct anatomical challenges compared to central subcortical targets. Here we investigate the region-specific artefacts encountered during activity and rest while streaming data from brainstem implants with a cranially-mounted device in two patients. Such artefacts result from this complex anatomical environment and its interactions with physiological parameters such as head movement and cardiac functions. The implications of the micromotion-induced artefacts, and potential mitigation, are then considered for future closed-loop stimulation methods.

## I. INTRODUCTION

Invasive central nervous system neuromodulation may be achieved through implantable devices, interfacing with sub-cortical structures through contact arrays of various configurations. In their most common clinically applied iteration, these are situated on linear leads, which are surgically inserted in the target area for stimulation (deep brain stimulation, DBS). DBS is a treatment method in a variety of pharmacoresistant neurological disorders, with subcortical targets ranging from clinical use in basal ganglia to the investigational exploration of the brainstem.

Novel neuromodulation methods utilise sensing capabilities in closed-loop strategies, where endogenous oscillations are targeted [1], [2]. Closed-loop designs are susceptible to artefact sensitivity however, which can be particularly prominent in areas where there is a physiological tendency for increased mobility. One such area with increased mobility due to its anatomical characteristics is the brainstem, a brain region with promising DBS targets for treatment of movement disorders, chronic pain, disorders of consciousness and sleep [3], [4].

The brainstem is encased in a system of spaces filled with cerebrospinal fluid (CSF), namely the quadrigeminal cistern, the cisterna pontis and interpeduncularis as well as the fourth ventricle towards its caudal end. Its extension, the medulla oblongata, is also dorsally surrounded by CSF through the cisterna cerebellomedullaris, while ventrally encased in the caudal portion of the cisterna interpeduncularis [5]. This surrounding CSF-filled network of spaces results in the brainstem being afforded a natural degree of motion during head movements on the midline and anterior-posterior axis. The rigid straight design of DBS electrode leads, in combination with local anatomical parameters and a higher brain compliance, may lead to micromotion of the implant. This phenomenon can lead to tissue strain, decreased interface longevity as well as artefacts impacting recording quality [6]–[8].

While studies of brainstem kinematics are mainly indicated in cases of pathology, such as Chiari malformations, cervical canal stenosis or instability at the craniocervical junction [9]–[12], brainstem micromotion has also been examined in physiological anatomy during general neuroanatomical studies. During head and neck flexion and extension, it has been noted that the pons normally deviates by 2 mm from the midline, as the clivo-vertebral angle changes from 9° to 22° [13], [14]. However, there are also reports that during flexion and extension the clivo-pontine distance did not statistically differ significantly at a group level in normal participants, despite a range of changes from 0.1–1.6 mm being reported [15].

Another source of displacement arising from the brainstem’s vicinity to CSF-filled spaces is that pulsatile fluid flows may cause additional motion of neural tissue. As dictated by the Monroe-Kelly doctrine, intracranial volume – comprised of CSF, parenchymal and intravessel (blood) volumes – is constant, therefore any change in the volume of one of its components requires compensation [16]. During arterial expansion following the cardiac systole, intravessel volume increases, thus CSF needs to be shunted out of the intracranial space following a downward motion through the spinal canal. Brainstem motion associated to cardiac systole may be relatively small in scale (0.1–0.5 mm) but occur periodically with each cardiac cycle [17]. Similarly, during coughing or the Valsalva manoeuver, increase in blood flow through volume transfer from chest vessels to the carotid system leads to CSF outflow changes and brief caudal then cranial displacement (around 2 mm) of the brainstem [18]. Both CSF volume and lymphatic flow change on a diurnal basis [19], [20], there is therefore an additional potential for brainstem positional shifts on a much longer timescale even where body position remains unchanged, as in the case of patients with a decreased level of consciousness and limited ability to mobilise.

Based on the above results and anatomical considerations, implants situated in the brainstem and following a trajectory parallel to its rostrocaudal axis will also have a degree of mobility following natural target micromotions. We set out to examine the impact of positional differences and cardiac activity on the quality of streamed local field potentials (LFPs) from two patients with brainstem DBS leads. Since the novel iteration of this device would be capable of closed-loop stimulation [21], we were specifically interested in artefacts generated by these simple physiological functions, given their implications on the accuracy and capabilities of the system.

## II. METHODS

### A. Surgical Approach and Targeting

The Pedunculopontine Nucleus (PPN) of the upper brainstem was bilaterally implanted, as part of a clinical trial examining autonomic and motor function modulation in multiple systems atrophy, a lethal neurodegenerative condition (MINDS NCT05197816). We used the Harvard Ascending Arousal Network Atlas [22] to visualise the location of the nucleus prior to electrode insertion. Details of the implantation technique can be found in [23], [24]. This method involves a frontal entry point and lead trajectory along the brainstem’s rostrocaudal axis (Fig. 1). Accurate lead placement was confirmed by fusing preoperative magnetic resonance imaging (MRI) and postoperative computed tomography (CT) scans.

**Fig. 1.**
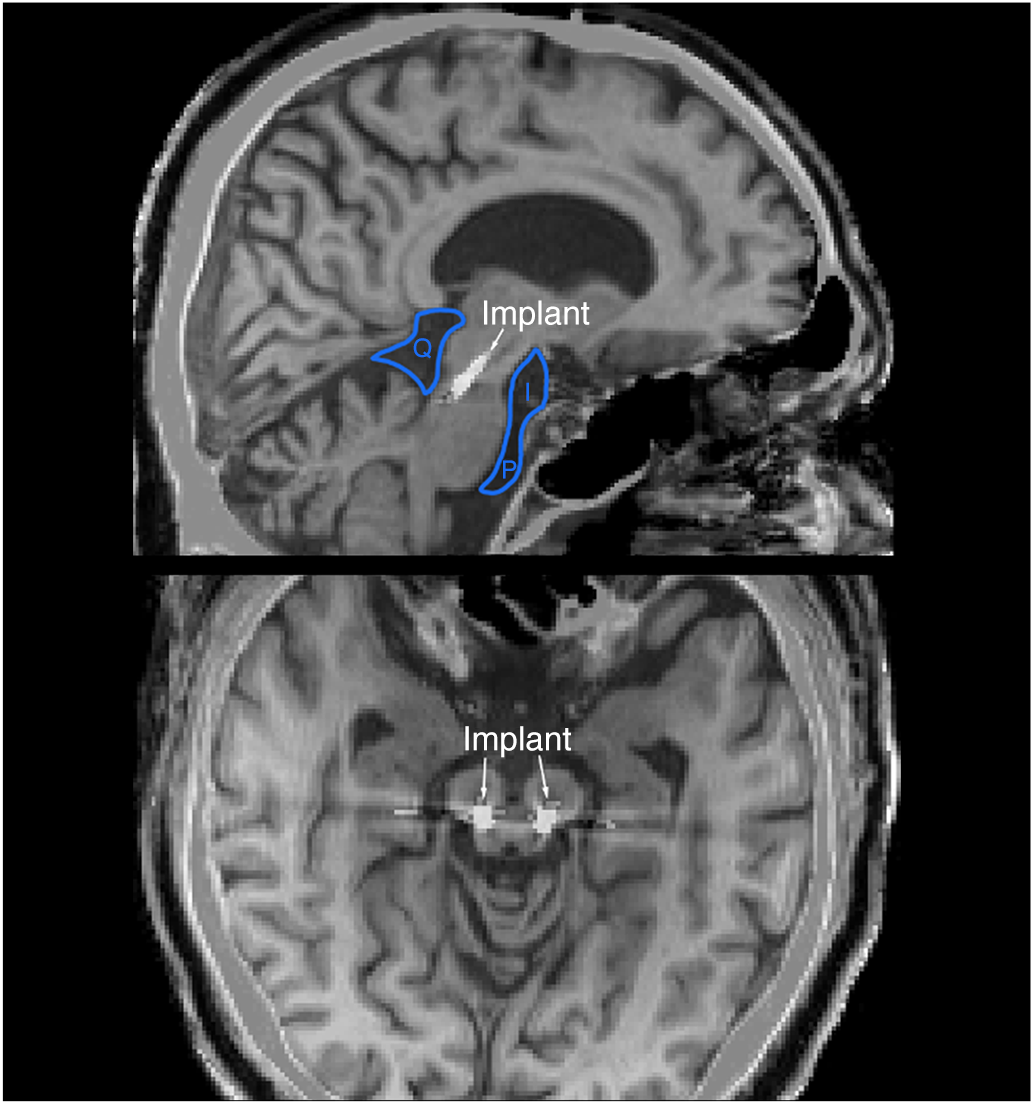
Placement of brainstem implants with relations to CSF-filled spaces. Saggital (top) and axial (bottom) views of the brain, with an implanted lead at the level of the PPN. I: Interpeduncular Cistern. P: Cisterna Pontis. Q: Quadrigeminal Cistern. Spaces filled with CSF (black) surround the implanted leads. Images generated by fusing postoperative CT scans with preopretaive T1-weighted MRI images for the same patient; metal appears white.

### B. Lead Specifications

We used DBS leads with four 1.5 mm annular contacts (with 1.5 mm spacing) (Bioinduction Ltd., Bristol, UK) as part of a cranially mounted system with LFP-streaming capability (at a sampling frequency of 1 kHz). The four contacts were sampled as series of dipoles, and located on a helically extruded lead body for optimal force distribution. These characteristics optimise lead viability by minimising shear stress in the implantation environment.

### C. Signal Streaming and Analyses

We streamed LFPs with simultaneous electroencephalogram (EEG), electromyogram (EMG) and electrocardiogram (ECG, time-locked to EEG and EMG). For motion artefact capture, we instructed patients to nod rhythmically, recording head motion artefacts on our extracranial electrode system, followed by a period of stable head position. Streaming was initiated prior to nodding and consisted of ‘sweeps’ of an average of 49.52 s (ranging 18.3–54.54 s).

For signal analyses, LFP files from two patients were loaded into Matlab (2022a, Mathworks Inc., Nantick, MA, USA), and pre-processed with a set of 3rd order Butterworth filters, a high-pass set at 0.1 Hz, and a low-pass at 180 Hz. For EEG, a band-stop for the rejection of 50 Hz line noise and its harmonics was also used. Dropped packages during the streaming process were accounted for and interpolated based on adjacent values. We did not initially de-trend the LFP signal, since we were interested in the presence of drifts as artefact. ECG was additionally processed with an adaptive thresholding operation and decision rules to identify QRS complexes and inter-beat intervals (IBI) [25], [26].

Both intracranial and extracranial signals were visually inspected to assess for common grossly visible artefacts and verification of arrhythmia versus noise presence in the processed ECG trace. Signals were time-locked based on common presence of QRS complexes and gross motion artefact, with results verified using known (marked) time lags between recording and streaming systems. Furthermore, frequencies of motion (as defined by extracranial electrode artefact) as well as cardiac pulsation (as defined by IBI during periods without arrhythmia) were calculated and subsequently defined as frequencies of interest (FOIs).

LFPs were further band-passed to artefact FOI, with power spectra visualised after a short-time Fourier transform (STFT, with a 5 s time window and 3 s overlap), then averaged across the FOI spectrum and compared across different lead contact dipoles. For constant cardiac artefacts, signal power within the artefact FOI was compared to power within the beta (15–30 Hz) and gamma (35–50 Hz) bands, as based on prior reports of human recordings these correspond to physiological PPN activity [4], [27]. Power within these activity bands was compared to pooled trials from both patients, for motion artefact-free periods averaged for the same dipole, using both pairwise t-tests with Bonferroni correction as well as a two-way ANOVA (with FOI and patient code as subgroup analyses) for pooled data.

With regards to motion, we noted that rhythmical head movements generated associated LFP artefacts (Fig. 2), which were not present while at rest. In addition, cardiac cycle-related artefacts were present in the signal (Fig. 3).

**Fig. 2.**
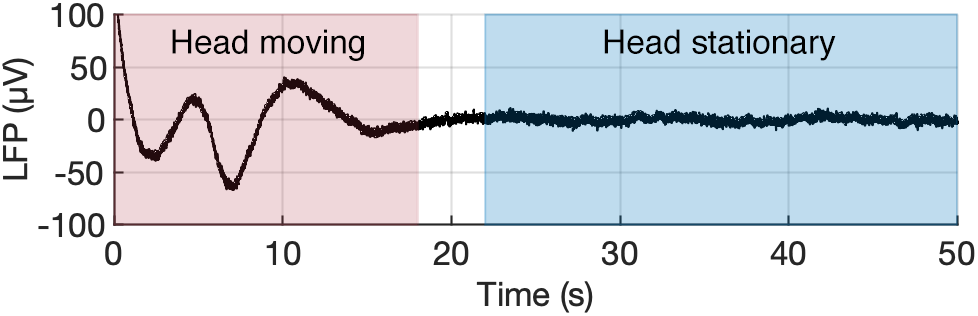
Periods of head motion, such as nodding (red) are characterised by a slow rhythmic artefact (< 0.5 Hz) in the streamed LFP (shown between top and bottom contact), absent while head position is stable (blue).

**Fig. 3.**
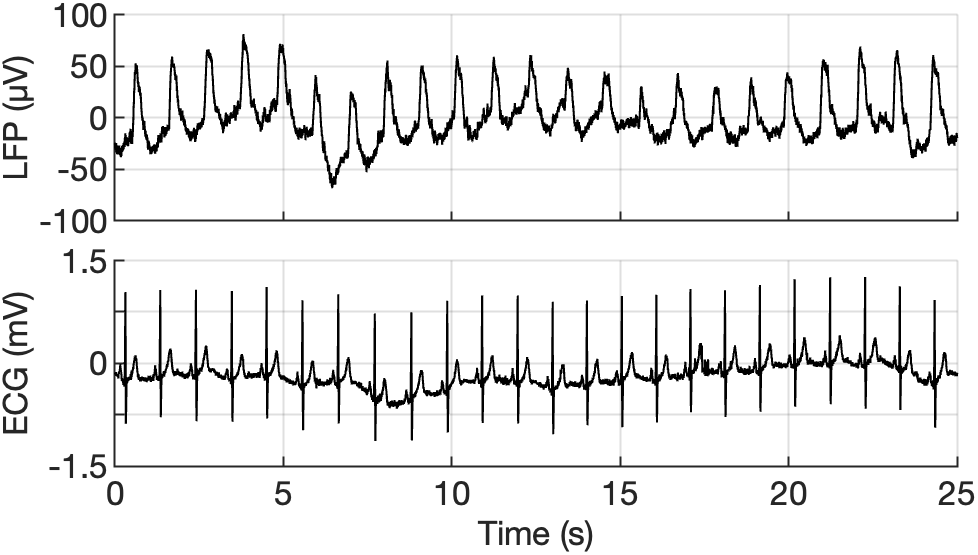
Prominent example of the periodic artefacts, in streamed LFP (between top and bottom contact) shown alongside a comparable period of ECG signal. Artefact peak frequency aligns with RR intervals.

Based on IBI after any ectopic heartbeat exclusion (and accounting for heart rate variability within patient) the artefact FOI was calculated, with upper limits that did not exceed 1.2 Hz and lower limit also affected by filtering properties (Fig. 4). We noted that the magnitude of the artefact varied between dipole configurations.

**Fig. 4.**
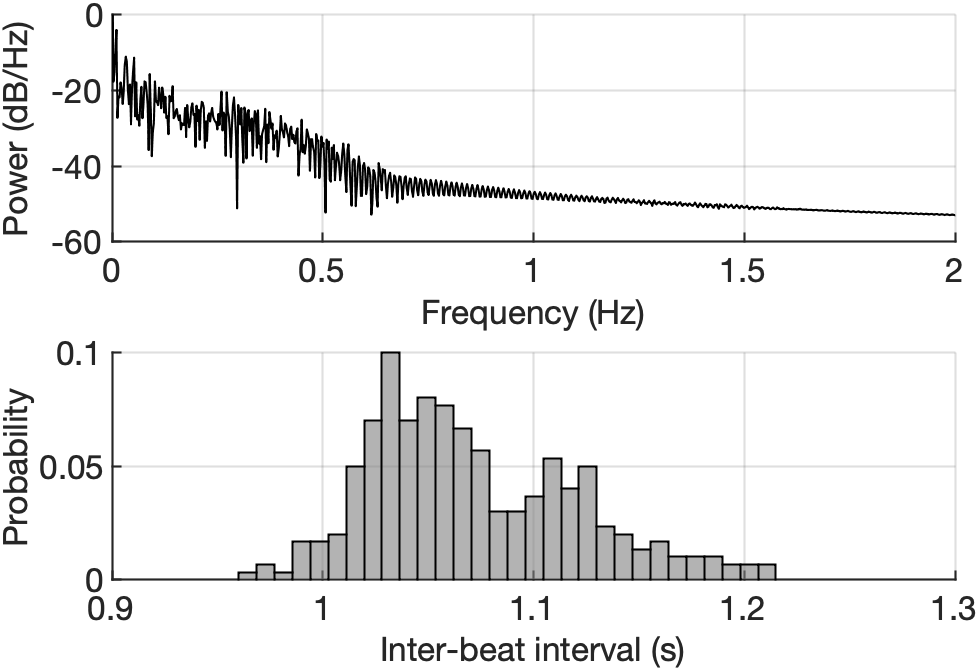
Characterisation of the Frequency of Interest (FOI) of the cardiac artifacts. Top: Welch spectral density of ECG signal. Bottom: distribution of IBIs over a 300 subsequent beats. Ectopic beats were excluded from the analysis.

More specifically, when caudal dipoles consisting of the adjacent lead contacts of two patients were compared between hemispheres, there was no significant difference between mean power in the IBI FOI (p = 0.16, CI = –1.63 to 9.55). When adjacent dipoles were compared to mid-lead contact dipole, which was less encased by a greater ratio of CSF to brain matter, the latter was more contaminated (p < 0.0001, CI = –39.18 to –30.63). Finally, when adjacent dipoles were compared to the largest potential dipole within the lead (top to bottom contacts), IBI FOIs differed significantly (p < 0.0001, CI = –60.50 to –44.75) with the largest streaming dipole being the most contaminated.

We subsequently explored the effects of cardiac artefacts on signal-to-noise ratio, especially during periods of optimal streamed signal quality without motion artefact. Mean power-in-band across all pooled dipole STFT time bins for the head stationary part of each sweep revealed that cardiac FOI is significantly larger compared to intrinsic frequencies (beta and gamma, as previously defined). Both patients exhibited the same phenomenon, without differences between cases (corrected p = 1.2, F = 3.17). Both within patient and across dipoles and time, power within the cardiac artefact FOI band dominated the signal (corrected p < 0.001 in all cases, F = 299.18 for patient one, F = 111.11 for patient two and F = 345.09 for pooled results). The artefact FOI was distinct from intrinsic activity frequency bands, which were still identifiable – albeit with lower mean power-in-band (Fig. 5).

**Fig. 5.**
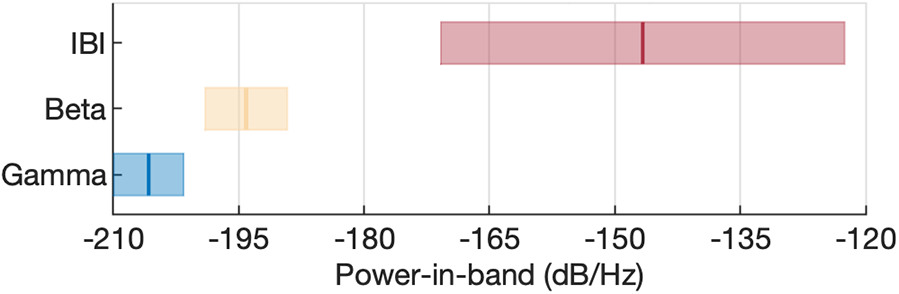
Mean power in different FOI bands, averaged across eleven 10 s stationary segments from each patient; bars represent one standard error. Despite cardiac artefact (IBI) dominating the power spectra, intrinsic PPN frequencies are still detectable.

## III. DISCUSSION

While the most frequent targets for invasive subcortical neuromodulation are located in the basal ganglia, brainstem nuclei are highly important in arousal regulation and therefore present an additional attractive target for intervention in disorders of sleep and consciousness. Previously, central thalamic stimulation showed promise in ameliorating behavioural impairments after severe traumatic brain injury [28], while brainstem targets could offer additional therapeutic benefits in cases of deafferentation with an intact extrathalamic pathway to cortex. While cardiac artefacts are expected to be present along ECG measurement axes within the body, prior modelling studies suggest that they should be largely absent in cranially mounted devices, in contrast to subclavicular devices [29]. This expectation has been reinforced by clinical results in patients with basal ganglia implants (subthalamic nucleus, external globus pallidus) [30]. However as we show, the brainstem’s unique anatomical complexity creates an additional pathway for cardiac artefacts to propagate, posing additional constraints to successful bi-directional device interfacing.

Bipolar re-referencing while acquiring and analysing signals from basal ganglia nuclei is an efficient strategy for artefact reduction [31], [32]; however, the relative velocity of micromotion during systole has been noted to be greater in the brainstem (by 0.5 mm/s) [33]. This could be due to the more extensive CSF encasement of the brainstem compared to basal ganglia – the only CSF-filled space adjacent to the subthalamic nucleus is the lateral ventricle in its rostral portion. The additional variance we noted (compared to basalganglia) highlights that region-specific anatomical constraints need to be carefully considered during signal analyses.

Specifically, we suggest that streaming and sampling from dipoles consisting of contacts which are unequally affected by CSF-distributed micromotion artefact should be discouraged except in cases of specific sub-regional interest, since the distribution of artefact FOI will not be eliminated from the signal. We showed that this phenomenon is present in brainstem sub-regions while at head rest due to artefact associated with the cardiac cycle (which therefore cannot be attenuated). However, in relatively stable dipoles consisting of adjacent contacts (thus maximising elimination of volume conduction) in regions further away from the pontine and quadrigeminal cisterns, the two hemispheres are not only comparable across patients but also minimally affected by low frequency artefacts.

Endogenous frequencies were still present in streamed LFP signals, albeit with a lower power-in-band, as endogenous PPN activity occurs mainly in the beta and gamma range, ensuring minimal overlap with cardiac IBI FOI. However, there are brainstem regions where tonic firing rates have been reported within that FOI, such as locus coeruleus (LC) baseline firing (around 1 Hz, increasing to 2 Hz in the presence of novelty in rodents) [34]. To our knowledge there is no reported case of microelectrode recordings in human LC, should this fact translate across species it could mean that meaningful signal is lost amongst IBI contamination. The position of LC is caudal-posterior meaning that conduction from the fourth ventricle would be a concern. However given its small diameter (mean length 14.5 mm, width 2.5 mm; height 2.0 mm [35], sampling translates to smaller inter-contact distance which could diminish common cardiac elements. Should brainstem stimulation become more widely applied, lead customisation could further facilitate its efficacy.

One of the main future challenges is the development of responsive stimulation, delivering state-specific therapy in accordance with endogenous brain activity. Such closed-loop systems of complex physiological states could be additionally optimised with the use of integrated extracranial activity sensing (such as motion detection through accelerometry). Such an additional sensor integration could lead to further artefact reduction in the case of brainstem targets, through gross motion artefact cancellation during streaming. In addition, while a closed-loop stimulation mode is active, an accelerometer could ensure safety and efficacy through a ‘fall-back’ mode activated by head motion. This could ensure state-specific stimulation (while immobile/inactive, to enhance arousal and performance) while avoiding sensing contaminants, especially in targets where the intrinsic activity frequency bands may be compromised during head motion or pose a risk of amplifier saturation.

Designing bioelectronics systems with advanced modulatory capabilities relies on identifying mitigations posed by physiological activity [36], and – as we show here – structural anatomy. Even in such a challenging micro-environment as the brainstem however, our investigation shows that close-loop responsive neurostimulation can be feasible, for instance through threshold setting on discrete endogenous oscillatory amplitudes. To further improve signal quality and eliminate the effects of cardiac artefact, additional steps could be implemented in the data acquisition algorithm, incorporating both feedback and adaptive control strategies [37]. The mean IBI can be approximated based on a regular ECG trace for each patient (routinely obtained during pre-operative screening), device sampling rates may then be adjusted accordingly, with interpolation of data to eliminate any artefacts associated with the QRS complex. However this would be based on an assumption that the mean IBI remains constant. Post-hoc deployment of an autoregressive model may further improve data quality, also taking into account gradual changes in mean heart rate but not ectopic beats, which may occur at unpredictable intervals [38]. In order to resolve both of these issues in vivo, an additional signal capturing ‘live’ heart rate would be required. Linking a peripheral sensor to the device, adjacent to a vessel that is located upstream from the DBS target (closer to the heart) would provide a brief window of opportunity to predictively eliminate the cardiac artefact, robust to ectopic beats. Further work in both device development and applied patient protocols will facilitate the creation of effective responsive systems, tailored to the requirements of this complex brain region.

## IV. LIMITATIONS

Our study is primarily limited by the small number of subjects and the specific patient pathology. Diurnal cycles of the CSF system [19] suggest future studies should account for time of day in the characterisation of artefact magnitudes.

## V. CONCLUSIONS

The introduction of closed-loop control policies presents significant potential for the refinement of neuromodulation; however, it relies on the accurate sensing of μV–scale physiological signals [21]. While the brainstem appears to be an attractive therapeutic target for disorders of consciousness [4], the presence of physiological artefacts poses a challenge to closed-loop algorithm development. Although the cardiac artefacts noted in this study could be mitigated through either software (e.g. template removal) or hardware (e.g. distributed sensing) solutions, these methods carry their own limitations. To ensure patient safety, risk assessments of closed-loop brainstem therapies should take this newly uncovered artefact class into account and provide fallback options in case of significant signal contamination.

## Acknowledgment

We thank Mr Sean Martin, Dr Sarangmat Naharaja and members of the Neurosciences Inpatient Service team for patient care and recruitment in the MINDS trial. We also thank the Bioinduction team for device-related guidance.

## Disclosures

The University of Oxford has research agreements with Bioinduction Ltd. Tim Denison also has business relationships with Bioinduction for research tool design and deployment, and stock ownership (< 1 %).

## Data Availability

The authors will consider requests to access the data that support the findings of this study in a trusted research environment. Contact: Alceste Deli.

